# Impairment of zebrafish reproduction upon exposure to melengestrol acetate

**DOI:** 10.1101/146506

**Authors:** Kewen Xiong, Chunyun Zhong, Xin Wang

## Abstract

Synthetic progestins contamination is common in the aquatic ecosystem, which may lead to serious health problem on aquatic animals. Melengestrol acetate (MGA) has been detected in the aquatic environment; however, its potential effects on fish reproduction are largely unclear. Here, we aimed to investigate the endocrine disruption and impact of MGA on zebrafish reproduction. Six-month old reproductive zebrafish were exposed to four nominal concentrations of MGA (1,10, 100 and 200 ng/L) for 15 days. Treatment with MGA reduced the egg production with a significant decrease at 200 ng/L. The circulating concentrations of estradiol and testosterone in female zebrafish or 11-keto testosterone in male zebrafish were significantly diminished compared to the non-exposed control fish. The early embryonic development or hatching rates were unaffected during the MGA exposure. Our results indicated that MGA was a potent endocrine disruptor in fish and the fish reproduction could be impaired even during a short-term exposure to MGA.

## Introduction

The normal endocrine functions of aquatic animals can be disturbed by the contaminants present in the aquatic environments [1, 2]. Most of the active compounds are detected in the environment are endocrine-disrupting chemicals, including natural and synthetic steroid [3–6]. Previous studies have reported the adverse effects of natural and synthetic steroidal estrogenson aquatic organisms, such as 17β-estradiol, 17α-estradiol, estrone, mestranol, ethinylestradiol [7–12]. However, little is known about the effects of MGA to the environmental health. Progestins have been widely used in veterinary medicine and hormonal therapies [13–15]. These chemicals have been found in sewage treatment plants, pharmaceutical industries and agricultural areas and are now world widely environmental pollution [16–18]. MGA is one of the most commonly used synthetic growth promoters, which is excreted in feces and urine of cattle [19]. MGA is an orally active progestagenic drug that has been used as a feed additive to beef cattle. It is primarily excreted from cattle unmodified and is very stable in soil and manure [19]. Several studies have reported changes in hormone concentrations, reproduction and morphology changes in aquatic organisms after exposure to trenbolone acetate or metabolites [20–26]. Studies investigating MGA effects in aquatic organisms are virtually unknown. Recently, natural progesterone or synthetic progestins have been reported that may result in endocrine disruption and impair reproduction in fish [12,18,27–32]. However, little is known about the potential underlying mechanisms. Even the synthetic progestins are widely used and detected in the environment; few researches have reported the hazards and their risk to the environment and the aquatic organisms. Currently, it is not known whether MGA had any effect on the reproduction in fish.

Therefore, we investigated the effect of MGA on the reproduction of zebrafish exposed to four nominal concentrations of MGA (1,10,100 and 200 ng/L). We also examined the sex hormone levels in zebrafish. The present study showed that MGA inhibited zebrafish reproduction in a dose-dependent manner after short-term treatment.

## Materials and methods

### Chemicals

Melengestrol acetate (MGA, 17α-Acetoxy-6-methyl-16-methylene-4,6-pregnadiene-3,20-dione, Cat. No. 33998) and DMSO (Dimethyl sulfoxide, Cat. No. D8418) were purchased from Sigma-Aldrich. MGA stock solution (1 mg/mL) was dissolved in DMSO and stored at −20 °C. All the other reagents used in this study were of analytical grade.

### Zebrafish husbandry

Six-month-old zebrafish AB line was maintained normally (temperature, 28 °C; pH 7.2-7.4; 14 hr on and 10 hr off light cycle).

### MGA treatment

The fish were housed in 50-L tanks with 30 L of water. Three replicate tanks containing 20 males and 20 females each were used. In order to obtain tank-specific baseline data for potential statistical comparison after initiation of chemical exposure, all the fish were maintained for a 14-day pre-exposure period. After the pre-exposure period and all the female fish have been successful spawned for several times, chemical treatment was then performed. The zebrafish were treated with different concentration of MGA (1,10,100 and 200 ng/L) or equal concentration of vehicle solution for 15 days, and the zebrafish embryos were collected daily. The water of the tanks exposed to MGA was changed daily. After 15 days of treatment with MGA, the eggs were collected and divided into two groups. One group was exposed to the same concentration of MGA (1,10,100 and 200 ng/L), and the other group received 0.001% (v/v) DMSO as the control.

### Tissue collection

After exposing to MGA for 15 days, the adult fish were anesthetized in 0.04% Tricaine (Sigma-Aldrich, Cat. No. A5040). Blood samples were collected from the caudal vein of the fish; the gonad were dissected and immediately stored at -80 °C for further analysis.

### Hormone measurement

Blood samples from three adult fish of the same gender were pooled, centrifuged (6,000 g for 10 min) at 4 °C to obtain the plasma. Plasma extraction and the measurement of the sex hormone levels were performed as described previously [33].

### Statistical analysis

All experiments were repeated three times independently. A one-way analysis of variance (ANOVA) with Tukey’s multiple comparisons was used to detect significant differences between the control and treated groups. Data were recorded as the mean with SD ± SE. A p <0.05 was considered statistically significant.

## Results

### Fish growth and survival

In the adult zebrafish groups, all the fish were survived after 15 days MGA treatment. MGA exposure had no obvious effect on the growth of adult fish, and on the hatching and malformation rates in the F1 embryos. All the hatching rates were over 90% and the malformation rates were lower than 5%.

### Reproduction

As shown in Fig. 1, embryos production was consistent and similar during the preexposure period among all groups. In the 15-days exposure period, female fish treated with 200 ng/L MGA spawned fewer eggs compared to the control group from day 10 to day 15. Other tested concentration of MGA showed slight inhibitory effect on the fish reproduction.

**Figure 1.**
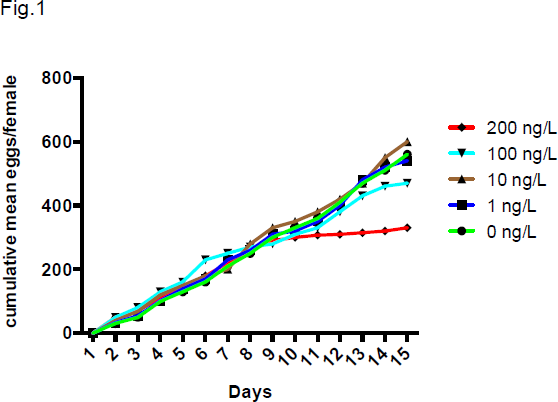
Effect of MGA exposure on the reproduction of female zebrafish.

### Sex hormone levels

In the female fish, treating with 10,100 and 200 ng/L of MGA significantly reduced the plasma E2 levies by 25%, 47% and 70%, respectively (Fig. 2A). Meanwhile, the testosterone levels were diminished by 15%, 22% and 30% in the 10,100 and 200 ng/L of MGA exposure groups, respectively (Fig. 2B). The plasma 11-KT levels were reduced by 42%, 55% and 61% after treatment with 10,100 and 200 ng/L of MGA, respectively (Fig. 2C).

**Figure 2.**
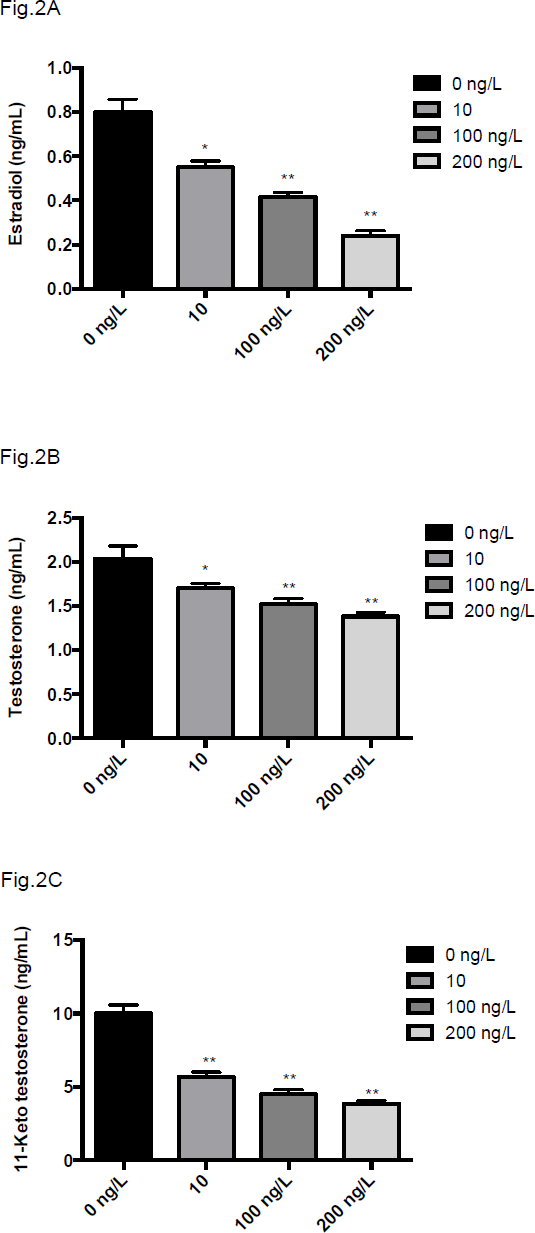
Effect of MGA treatment on plasma concentrations of E2 (A), testosterone (B) in female fish and 11-ketotestosterone (C) in male fish.

## Discussion

The synthetic progestins have recently been reported that may result in endocrine disruption and inhibit reproduction in fish [18, 27, 34, 35]. However, the toxicological effects and mechanisms of the synthetic progestin MGA on fish have not been evaluated yet. In this study, we found that the zebrafish reproduction could be impaired when exposing to MGA at the relevant environmental concentrations. The 11-KT levels in male zebrafish and the E2 and testosterone levels in female zebrafish were significantly diminished in MGA treated groups. There were no significant differences in embryos hatching, survival rate and developmental malformation during the exposure. In fish, one of the key ecologically indicators of endocrine disruption is the embryos production in females [36]. MGA exposure caused significant decrease in the egg production, which has not been reported till date. Recently, similar effects of MTA and EE2 with different combinations of progestins on fish fecundity have been reported [37, 38]. Hence, the potential risks of synthetic hormone to the fish species should be highlighted. Future studies are required to investigate the potential mechanisms of the hormonal effects on fish fecundity.

